# Baroreceptor denervation reduces inflammatory status and worsens cardiovascular collapse during systemic inflammation

**DOI:** 10.1101/748608

**Authors:** Mateus R. Amorim, Júnia L. de Deus, Camila A. Pereira, Luiz E. V. da Silva, Gabriela S. Borges, Nathanne S. Ferreira, Marcelo E. Batalhão, José Antunes-Rodrigues, Evelin C. Cárnio, Rita C. Tostes, Luiz G. S. Branco

**Author notes:** Corresponding author at: Dental School of Ribeirão Preto, University of São Paulo, Avenida do Café s/n, University of São Paulo, 14040-904 Ribeirão Preto, SP, Brazil. E-mail address (L.G.S. Branco) (M.R. Amorim). **Funding sources:** LGSB [#2016/17681-9, São Paulo Research Foundation (FAPESP)], MRA [#2017/09878-0 São Paulo Research Foundation (FAPESP)]; National Council for Scientific and Technological Development (CNPq), Brazil.

## Abstract

Beyond the regulation of cardiovascular function, baroreceptor afferents play polymodal roles. We hypothesized that baroreceptor denervation affects lipopolysaccharide (LPS)-induced systemic inflammation (SI) and hemodynamic collapse in conscious rats, and that these parameters are interconnected. We combine: a) hemodynamic and thermoregulatory recordings after LPS administration at a septic-like dose b) analysis of the cardiovascular complexity, c) evaluation of vascular function in mesenteric resistance vessels, and d) measurements of inflammatory cytokines (plasma and spleen). LPS-induced drop in blood pressure was higher in sino-aortic denervated (SAD) rats. LPS-induced hemodynamic collapse was associated with SAD-dependent autonomic disbalance. LPS-induced vascular dysfunction was not affected by SAD. Surprisingly, SAD blunted LPS-induced surges of plasma and spleen cytokines. These data indicate that sino-aortic afferents are key to alleviate LPS-induced cardiovascular collapse, affecting the autonomic cardiovascular control, without affecting resistance blood vessels. Moreover, baroreflex modulation of the LPS-induced SI and hemodynamic collapse seem not to be interconnected.

## Introduction

Sepsis is a common disorder affecting 31.5 million people worldwide, with a mortality rate of 5.3 million deaths every year (1). Considering the severity of sepsis and its pathophysiological complications, different research groups have focused on the study of mechanisms of systemic inflammation (SI) in search of therapeutic strategies help manage signs and symptoms for the treatment of this condition (2–4). Systemic administration of lipopolysaccharide (LPS) has been widely used in animal models to induce changes observed during SI, such as exacerbated production and release of cytokines, catecholamines, hormones and nitric oxide (NO), associated with hypotension, tachycardia, and hypothermia followed by fever (5–9). These hemodynamic and thermoregulatory responses to LPS are similar to those observed in humans with sepsis (10).

Classically, baroreceptor afferents are specialized structures in the homeostatic control of blood pressure in health and disease (11, 12). Baroreceptors afferents integrity is mandatory in the control of blood pressure within a narrow range of variation and surgically removed baroreceptors afferents rats [sino-aortic denervation (SAD)] had higher variability of the mean arterial pressure (11,13,14). More recently, it has been shown the role of baroreceptor afferents mediating LPS-induced cardiovascular collapse (15–18). Moreover, in SI, electrical baroreflex stimulation in rats has been reported to blunt LPS-induced production of cytokines in the hypothalamus (19), indicating an anti-inflammatory role played by aortic depressor nerve stimulation. However, the involvement of the baroreceptors afferents in the inflammatory status has not received the same attention, neither its putative relation with the cardiovascular system.

In this study we examined in an integrated matter to the role of baroreceptor afferents integrity in LPS-induced classical inflammatory cytokines surges [in plasma and spleen – effector organ of the splenic anti-inflammatory reflex (2, 20)], hypothermia and fever, as well as the relation of the inflammatory status with cardiovascular function in rats.

## Results

### Cardiovascular and thermoregulatory changes during LPS-induced SI is dependent on sino-aortic afferents integrity

First, we investigated whether LPS-induced responses in mean arterial pressure (MAP) and heart rate (HR) were affected by the surgical removal of the arterial baroreceptors (SAD). MAP (P = 0.5189) and HR (P = 0.2197) were similar in Control + Sal and SAD + Sal animals throughout 180 min (Fig. 1 A). LPS-induced fall in MAP was significantly higher (P < 0.0001) and occurred earlier (P = 0.0413) in SAD + LPS in comparison to Control + LPS rats (Fig. 1 B and D). Furthermore, LPS-induced tachycardia was blunted in SAD + LPS in comparison to Control + LPS rats (Fig 1 B, P < 0.0001). Interestingly, SAD rats presented no fever, but a significant drop in Tb (hypothermia) in response to LPS (Fig. 1 F and G, P = 0.0075). These data indicate that hemodynamic and thermoregulatory control during LPS-induced SI depends on the sino-aortic afferents integrity. Successfulness of SAD surgery was confirmed using pharmacological activation of the baroreflex with phenylephrine (Phe, Fig. 1 C and E, P < 0.0001).

**Fig. 1.**
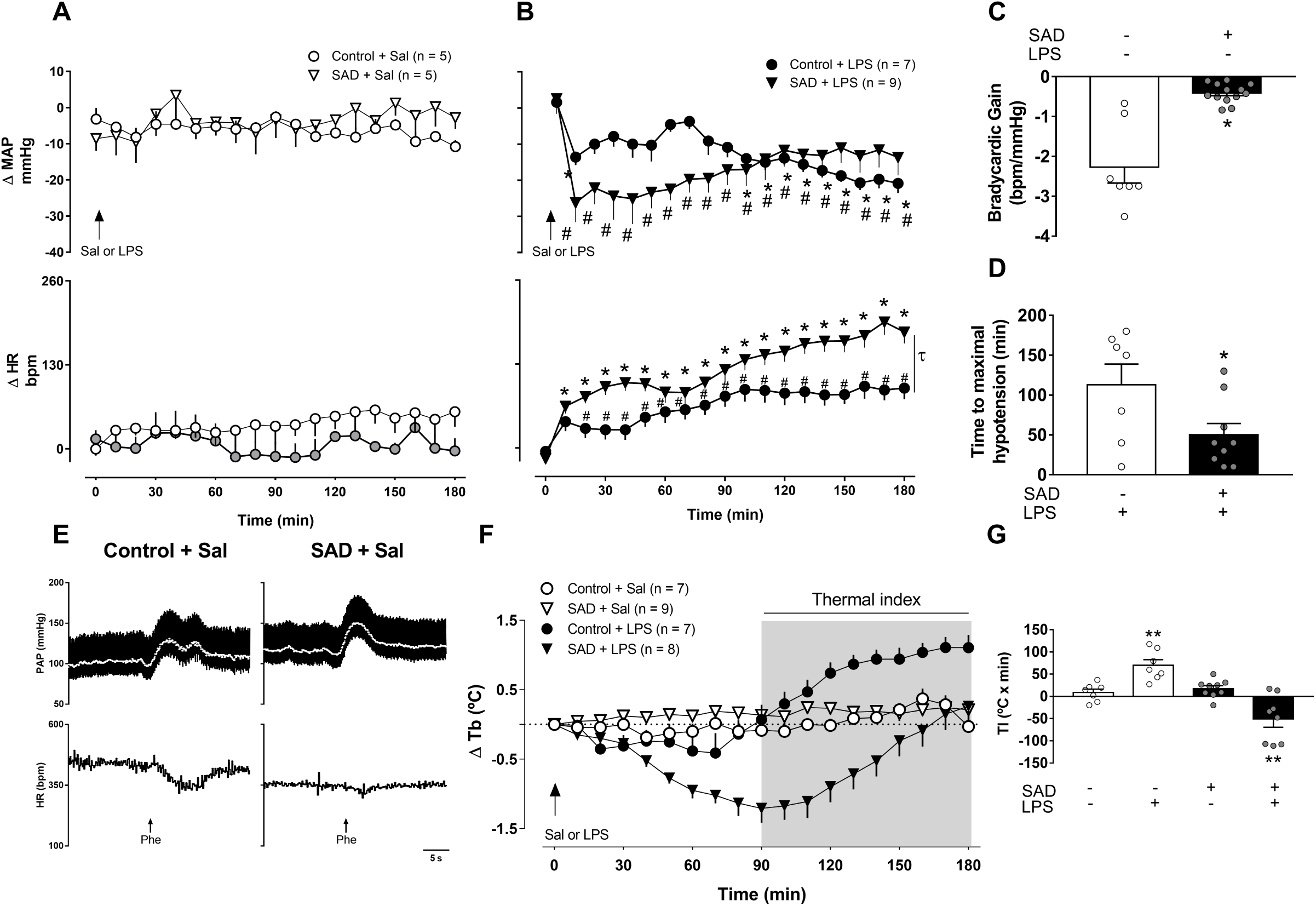
Effects of sino-aortic denervation (SAD) on mean arterial pressure (MAP), and heart rate (HR) of saline (Sal)-treated rats (panel A) and rats with SI (panel B, LPS, 1.5 mg.kg^−1^). Bradycardic gain during stimulation of the arterial baroreceptors (panel C) and maximal hypotensive response to LPS-induced SI (panel D). Representative recordings showing PAP, MAP (white line; top) and HR responses to i.v. injection of phenylephrine (Phe; panel E). Δ Tb (panel F) and thermal indexes (panel G). Results are presented as individual values and mean ± SEM. *, ^#^ p < 0.05 compared with time zero and τ p < 0.0001 difference between groups using the two-way ANOVA with Bonferroni’s post hoc test (panel A and B). *p < 0.05 using the unpaired t test (panel C and D). **p < 0.01 compared with Control + Sal group using the one-way ANOVA with Bonferroni’s post hoc test (panel G). Control + Sal (*n* = 5-7), Control + LPS (*n* = 7-9), SAD + Sal (*n* = 5-9), and SAD + LPS (*n* = 8-9).

### Variability in HR and SAP during LPS-induced SI

Variability of pulse interval (PI) in the time domain, determined by the standard deviation of normal to normal PI (SDNN) and root mean square of successive differences (RMSSD), was significantly reduced in Control + LPS (P < 0.0001 and P = 0.0004) and SAD + LPS (P < 0.0001 and P = 0.0009) in comparison with Control + Sal (Fig. 2 A and B). Variability of SAP in the time domain [evaluated by standard deviation (SD) of SAP] was significantly increased in SAD + Sal (P = 0.0041), but not in SAD + LPS (P > 0.9999) in comparison with Control + Sal animals (Fig. 2 C).

**Fig. 2.**
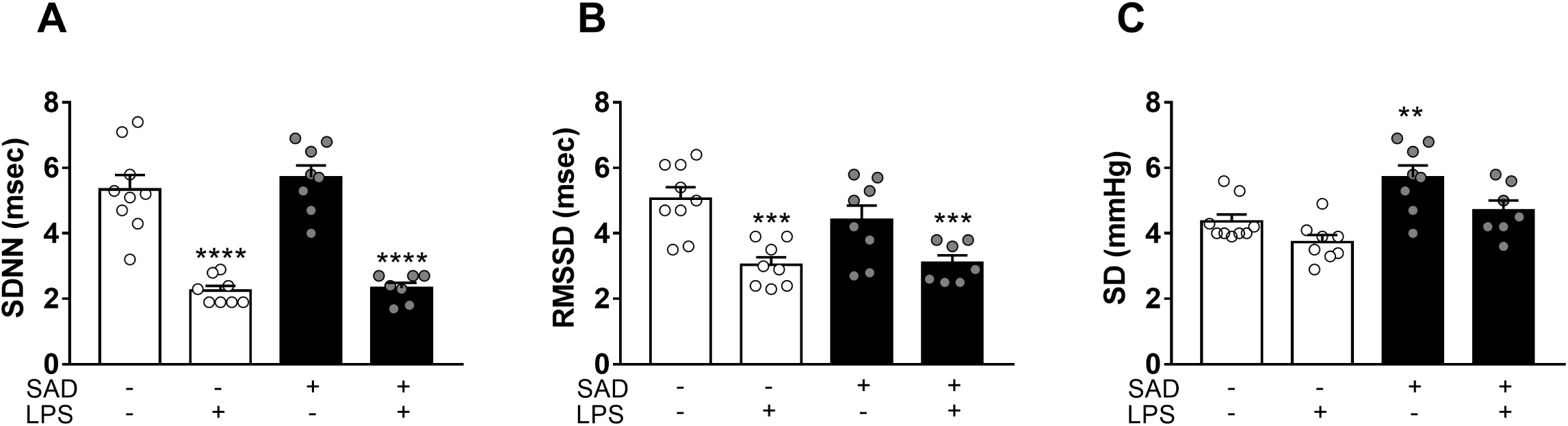
Variability in heart rate (HR) and in systolic arterial pressure (SAP) in the time-domain during LPS-induced SI. Standard deviation (SDNN, panel A) and root mean square of the successive differences (RMSSD, panel B) from PI series. Standard deviation (SD) from SAP series (panel C). Control + Sal, Control + LPS, SAD + Sal, and SAD + LPS groups were evaluated 3 and 24 hours after LPS administration. Results are presented as individual values and mean ± SEM. ** p < 0.01, *** p < 0,001, **** p < 0.0001 compared with Control + Sal group using the one-way ANOVA with Bonferroni’s post hoc test. Control + Sal (*n* = 9), Control + LPS (*n* = 8), SAD + Sal (*n* = 9), and SAD + LPS (*n* = 7).

Spectral analysis in the frequency domain of the PI showed that LF power was not significantly altered between Control + Sal, Control + LPS, SAD + Sal and SAD + LPS groups (Fig. 3 A and C, P > 0.9999). Otherwise, HF power was significantly reduced in Control + LPS (P < 0.0001) and in SAD + LPS rats (P < 0.0001) in relation to Control + Sal group (Fig. 3 B and D). These results indicate that LPS administration decreases cardiac vagal modulation in Control and SAD rats. Considering the SAP spectral analysis, a significant increase in the LF component was observed in the Control + LPS (P = 0.0051), but not in SAD + LPS (P > 0.9999) in relation to Control + Sal group, indicating that LPS-induced increase in the sympathetic vasomotor modulation depends on the baroreceptors afferents integrity (Fig. 3 B and E).

**Fig. 3:**
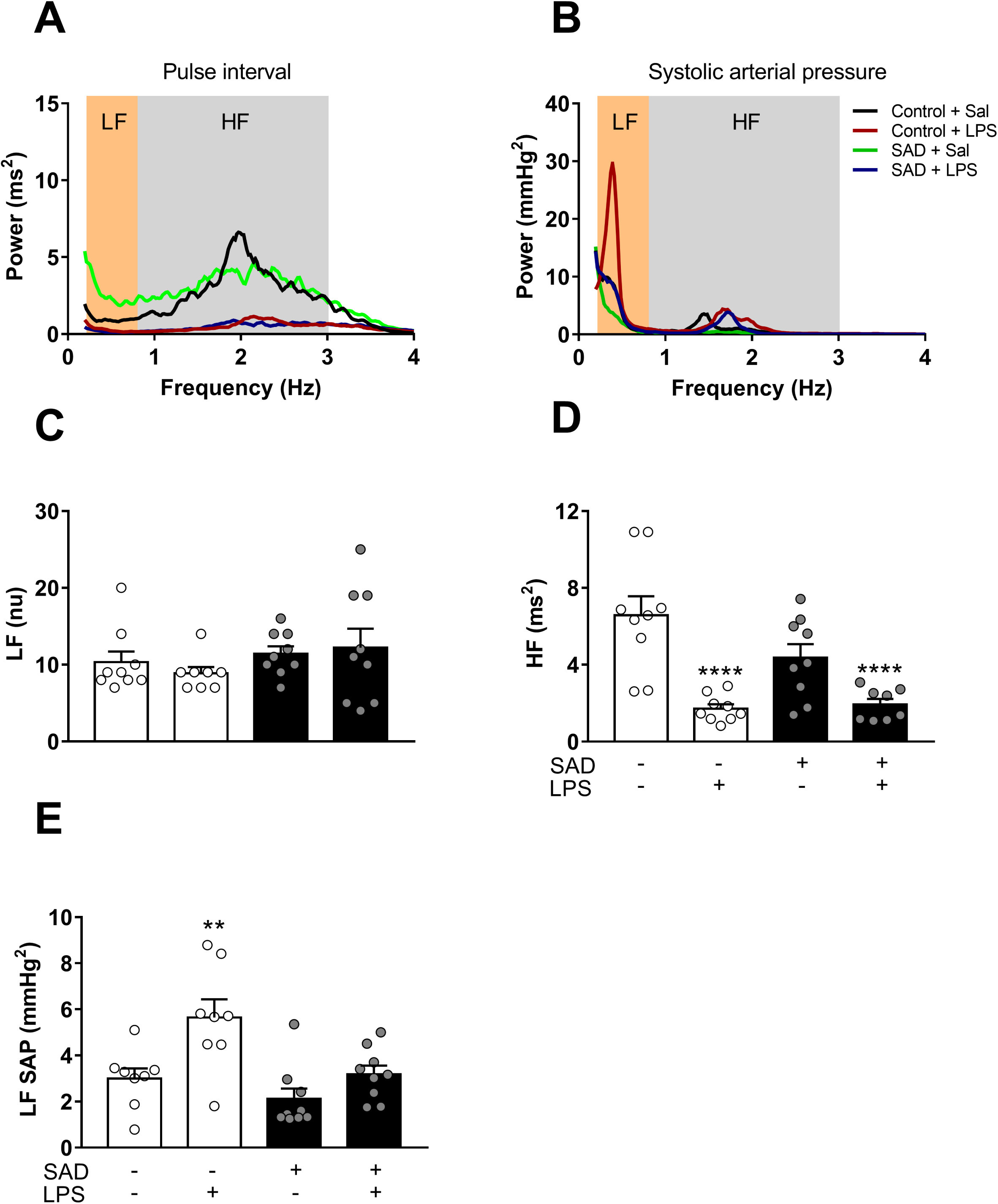
Power spectral analyses of pulse interval (PI) and systolic arterial pressure (SAP) from Control + Sal, Control + LPS, SAD + Sal, and SAD + LPS groups. Representative tracings from each experimental group (panel A and B). Magnitude of low frequency (LF, panel C) and high frequency (HF, panel D) components of PI. Magnitude of HF component of LF component of PI (panel E). Results are presented as individual values and mean ± SEM. ** p < 0.01, and **** p < 0.0001 compared with Control + Sal group using the one-way ANOVA with Bonferroni’s post hoc test. Control + Sal (*n* = 8-9), Control + LPS (*n* = 8-9), SAD + Sal (*n* = 8-9), and SAD + LPS (*n* = 8-9).

The detrended fluctuation analysis (DFA α_2) scaling exponent was lower in Control + LPS than in the Control + Sal group (P = 0.0223, Fig. 4 B), whereas the same scaling exponent was significantly increased in SAD + Sal (P < 0.0001) and in SAD + LPS (P = 0.0040) in relation to Control + Sal rats (Fig. 4 B). The DFA α_3 scaling exponent was significantly higher in SAD + Sal (P = 0.0133) than in Control + Sal rats (Fig. 4 C). The multiscale entropy (MSE) curves from all the evaluated groups are shown in Fig. 5 A and B. The MSE for the SAD + Sal group was significantly reduced on small time scales in relation to Control + Sal (P < 0.0001; P = 0.0004 and P = 0.0017; Fig. 5 A, C, D and E). On the other hand, in the Control + LPS, the MSE was significantly increased in comparison with Control + Sal group (P = 0.0444, Fig. 5 B and B).

**Fig. 4:**
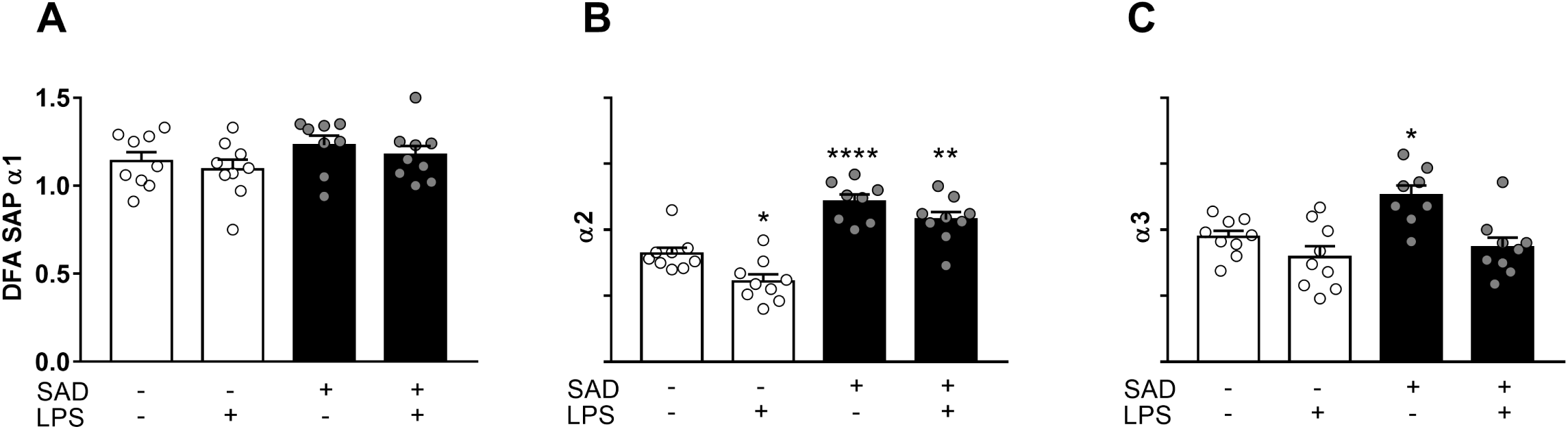
Detrended fluctuation analysis (DFA) from Control + Sal, Control + LPS, SAD + Sal, and SAD + LPS groups that where evaluated 3 hours after LPS administration. Average values of α1 (panel A), α2 (panel B) and α3 (panel C) are shown. Results are presented as individual values and mean ± SEM. * p < 0.05, ** p < 0.01, **** p < 0.0001 compared with Control + Sal group using the one-way ANOVA with Bonferroni’s post hoc test. Control + Sal (*n* = 9), Control + LPS (*n* = 9), SAD + Sal (*n* = 8), and SAD + LPS (*n* = 9).

**Fig. 5:**
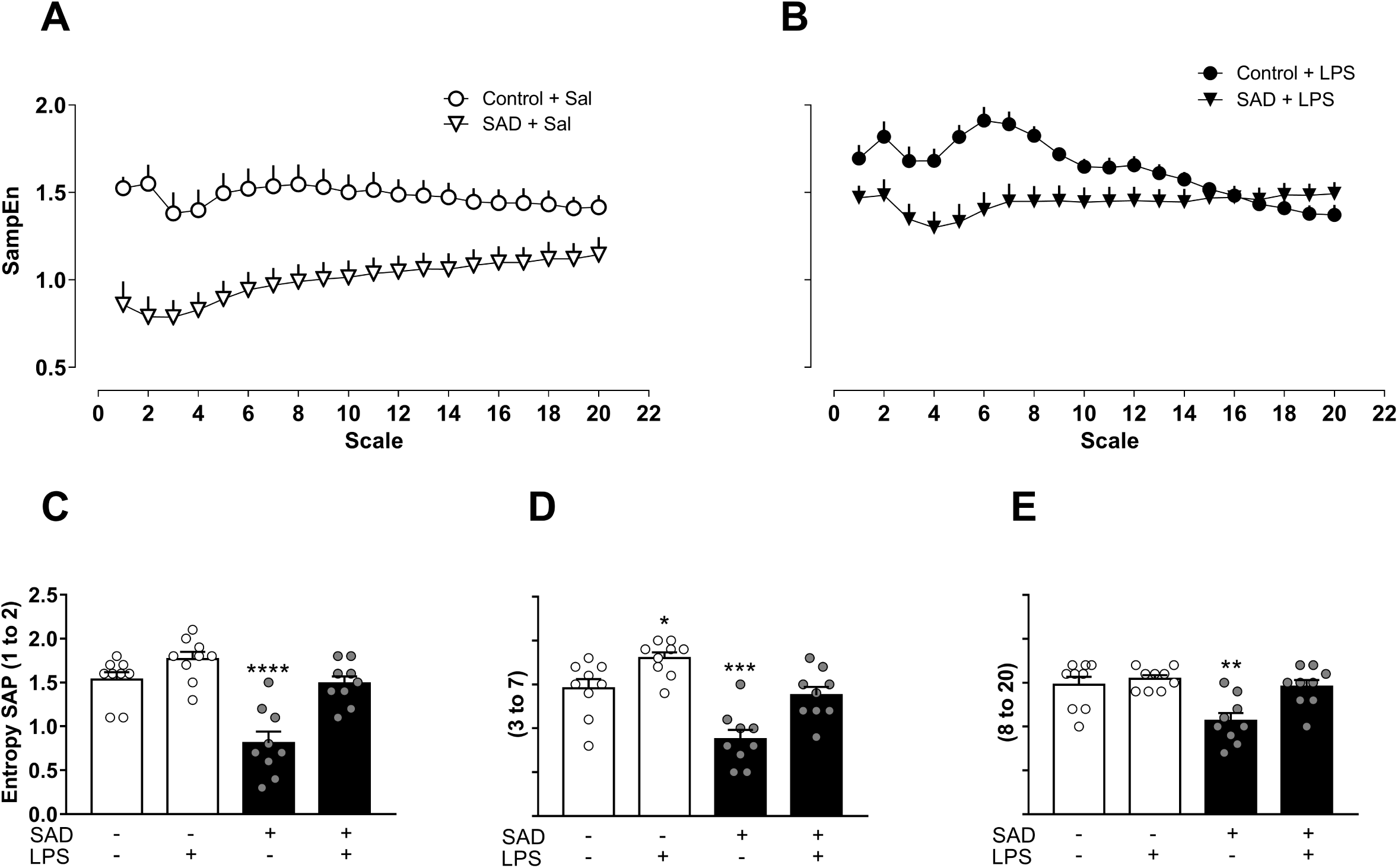
Multiscale entropy (MSE) from Control + Sal, Control + LPS, SAD + Sal and, SAD + LPS groups that where evaluated 3 hours after LPS administration. Mean MSE profiles obtained from Control + Sal, SAD + Sal (panel A), Control + LPS, and SAD + LPS (panel B). Average values of entropy calculated for scales 1 and 2, 3 to 7 and 8 to 20 are shown (panel C). Results are presented as individual values and mean ± SEM. * p < 0.05, ** p < 0.01, *** p < 0,001, **** p < 0.0001 compared with Control + Sal group using the one-way ANOVA with Bonferroni’s post hoc test. Control + Sal (*n* = 9), Control + LPS (*n* = 9), SAD + Sal (*n* = 9), and SAD + LPS (*n* = 9).

### Vascular reactivity during LPS-induced SI

Considering that LPS induced a significant drop in MAP associated with autonomic dysfunction, we investigated whether SI leads to vascular damage in resistance blood vessels and if this eventual vascular dysfunction is exacerbated in rats submitted to SAD. The contraction of mesenteric resistance arteries stimulated by potassium chloride (KCl - 120 mM) was significantly reduced in Control + LPS, SAD + Sal, and SAD + LPS when compared with Control + Sal group (P = 0.05, Fig. 6 A). Similarly, maximum contractile responses induced by Phe were reduced in mesenteric arteries from LPS-treated Control and SAD groups (P < 0.05, Fig. 6 B). Vascular hyporesponsiveness to vasoconstrcitors was reverted by the incubation with a non-selective inhibitor of nitric oxide synthase, L-NAME (P > 0.05, Fig. 6 C). These data indicate that both SAD and LPS administration *per se* leads to vascular dysfunction, but without additive effects.

**Fig. 6:**
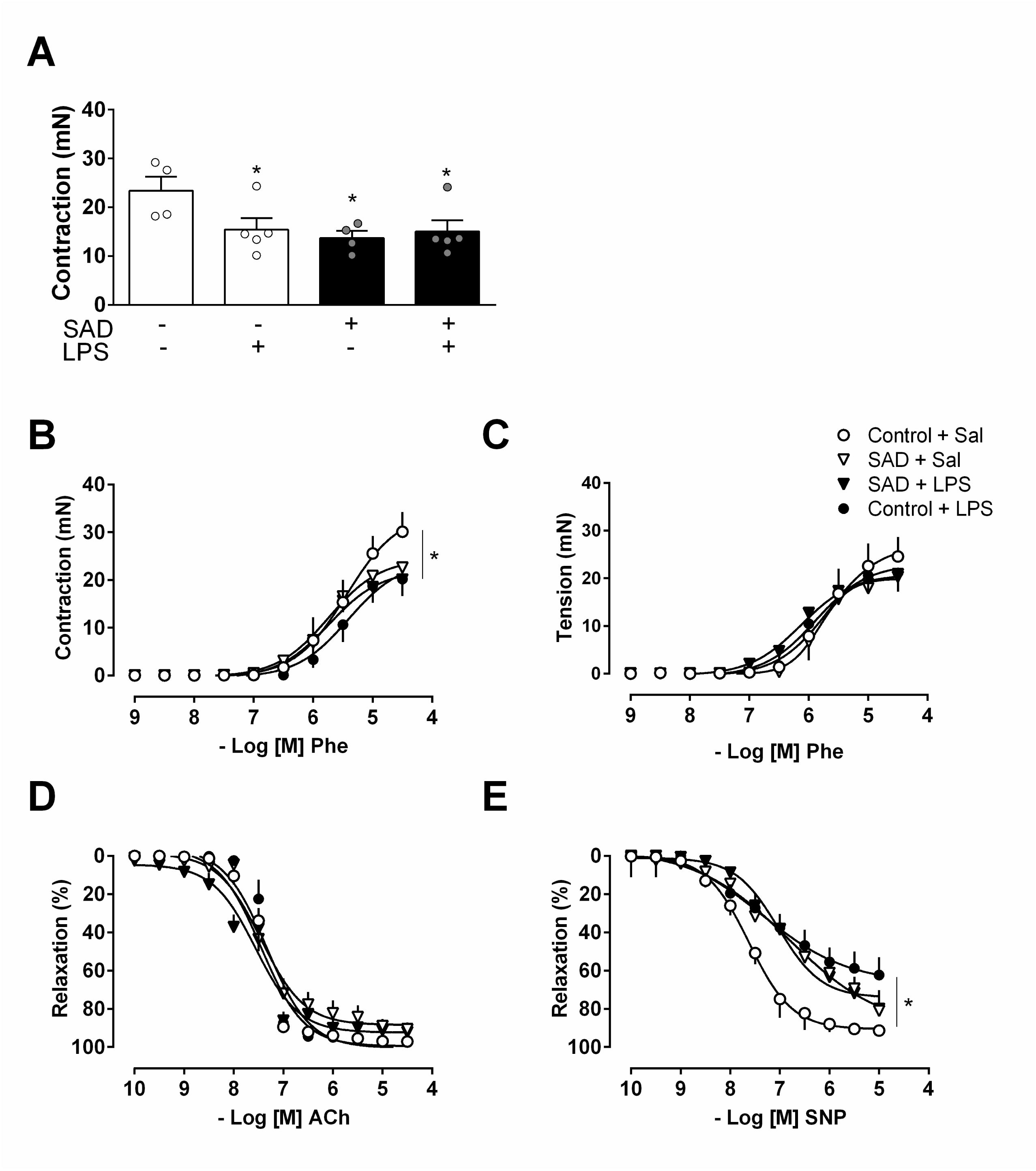
Vasoconstrictor responses to KCl (120 mM, panel A) from Control + Sal, Control + LPS, SAD + Sal and, SAD + LPS groups. Cumulative concentration-response curves to the α-1 adrenergic agonist [phenylephrine (Phe) panel B], Phe in presence of L-NAME, non-selective inhibitor of nitric oxide synthase, (10^−4^ M, panel C), Acetylcholine (ACh), endothelium-dependent vasodilator (panel D), sodium nitroprusside (SNP) endothelium-independent vasodilator (panel E). * p < 0.05 compared with Control + Sal group using the one-way ANOVA with Bonferroni’s post hoc test. Control + Sal (*n* = 4-5), Control + LPS (*n* = 5), SAD + Sal (*n* = 4), and SAD + LPS (*n* = 5).

Furthermore, endothelium-dependent vascular relaxation induced by cumulative concentrations of acetylcholine (ACh) was similar among the groups (Fig. 6 D). In contrast, endothelium-independent vasodilation to sodium nitroprusside (SNP) was reduced in mesenteric arteries from Control + LPS, SAD + Sal, SAD + LPS groups in comparison with the Control + Sal group (P < 0.05, Fig. 6 E). In addition, maximum contractile responses induced by electrical-field stimulation (EFS) were significantly decreased in arteries from Control + LPS, SAD + Sal and SAD + LPS rats (P < 0.05, Fig. 7 A). L-NAME reversed decreased EFS-induced contractions only in arteries from the Control + LPS group (P < 0.05, Fig. 7 B).

**Fig. 7:**
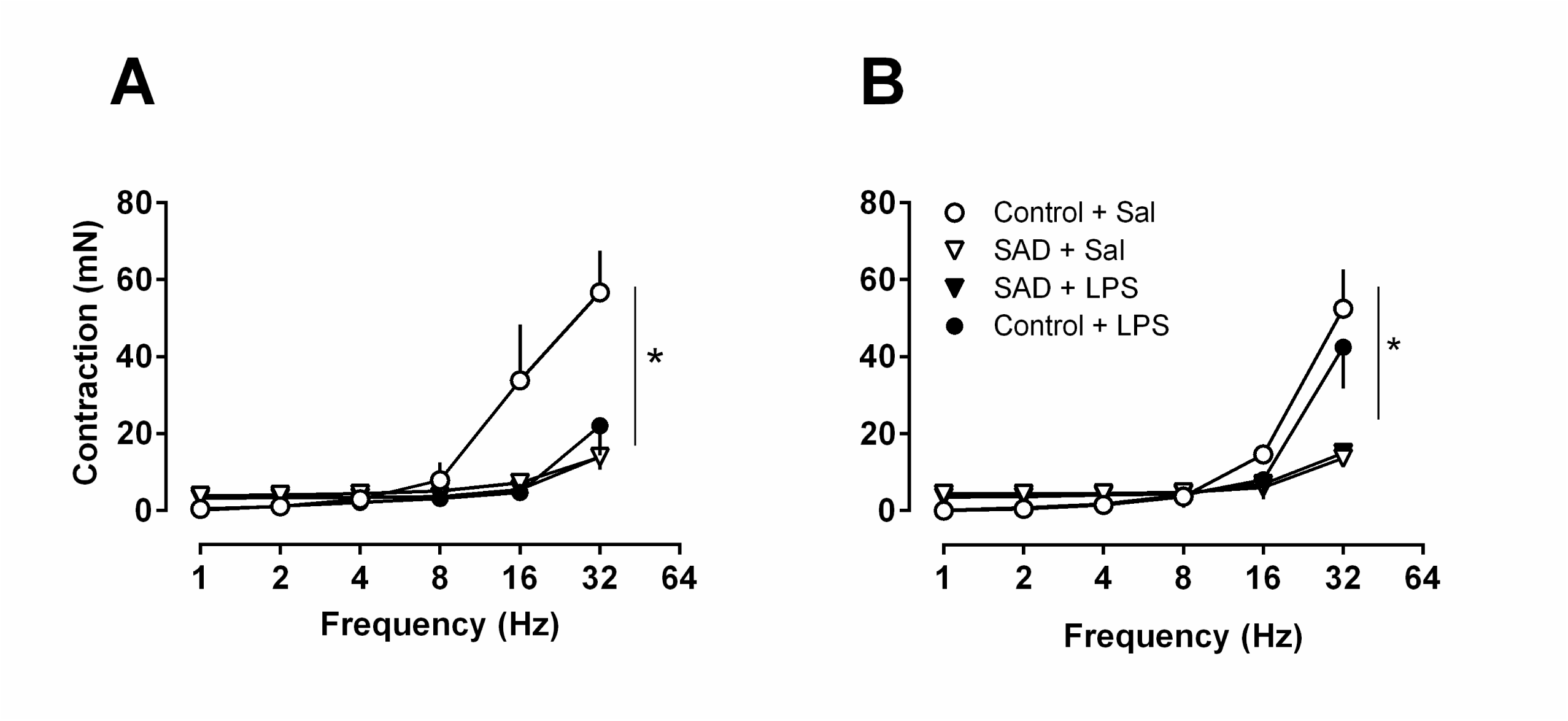
Electrical-field stimulation (EFS, panel A) from Control + Sal, Control + LPS, SAD + Sal and, SAD + LPS groups. EFS in presence of L-NAME (10^−4^ M, panel F) in resistance mesenteric arteries. Results are presented as mean ± SEM * p < 0.05 compared with Control + Sal group using the one-way ANOVA with Bonferroni’s post hoc test. Control + Sal (*n* = 4-5), Control + LPS (*n* = 5), SAD + Sal (*n* = 4), and SAD + LPS (*n* = 5).

### Cytokine levels in plasma and spleen during LPS-induced SI

Considering our previous study that documented hemodynamic and inflammatory changes 3 hours following LPS administration (18), cytokine levels were evaluated at this same period in plasma and spleen as an index of SI and the modulatory role of sino-aortic afferents on the splenic anti-inflammatory reflex, respectively.

#### Plasma

LPS increased the plasmatic levels of the pro-inflammatory cytokines TNF-α (P = 0.0030), IL-6 (P < 0.0001), and IFN-γ (P = 0.0086) and the anti-inflammatory cytokine IL-10 (P = 0.0002) in Control + LPS in comparison with Control + Sal rats. Interestingly, SAD decreased LPS-induced IL-6 (P = 0.0049) and IL-10 plasma levels (P = 0.05, Fig. 8 A, B, C and D). These results indicate that baroreflex positively modulates LPS-induced peripheral cytokine surges.

**Fig. 8:**
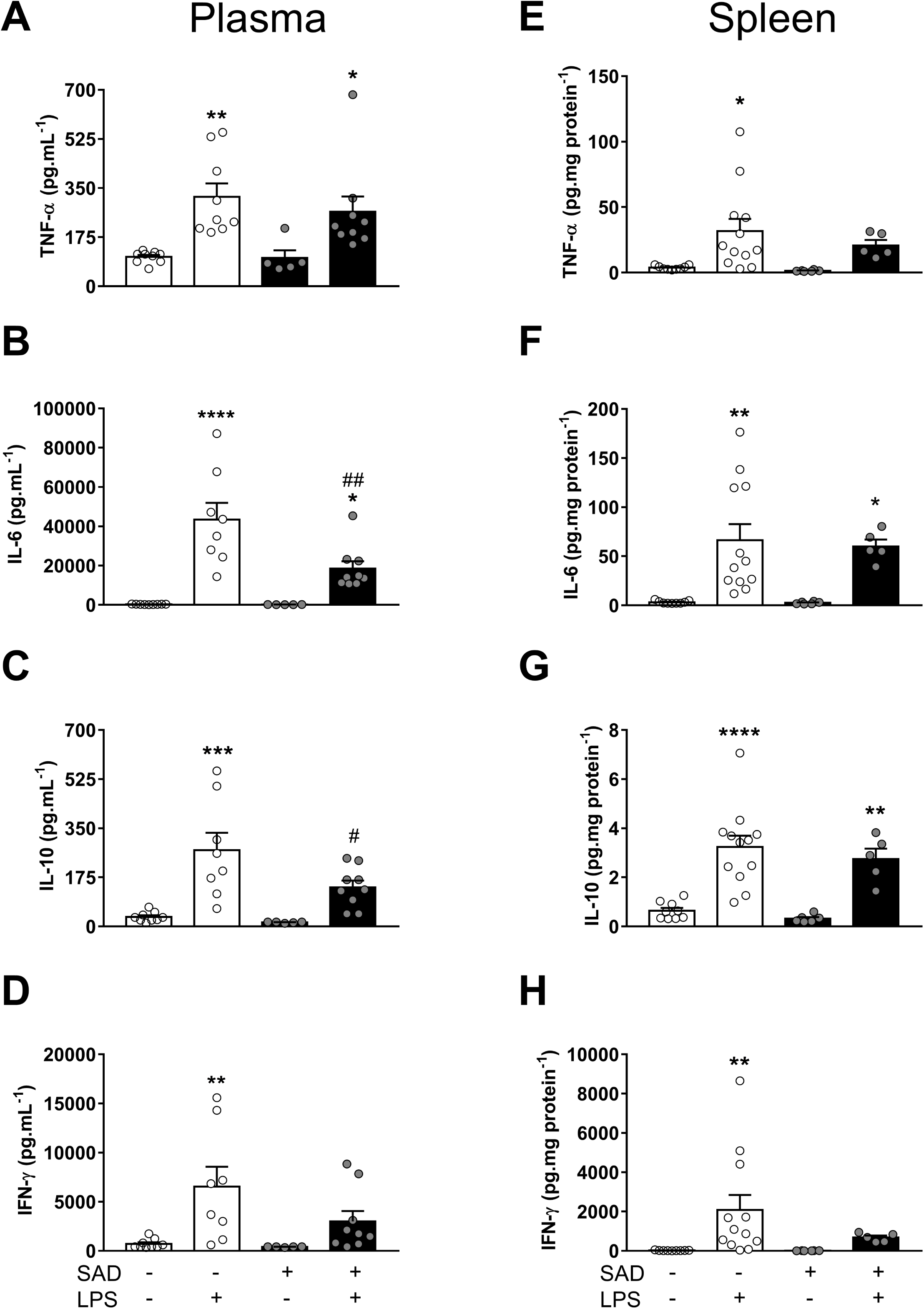
Plasma (panel A, B, C, and D) and splenic levels (E, F, G, and H) of pro-inflammatory and anti-inflammatory cytokines from Control + Sal, Control + LPS, SAD + Sal and SAD + LPS groups. Results are presented as individual values and mean ± SEM. * p < 0.05, ** p < 0.01, *** p < 0,001, **** p < 0.0001 compared with Control + Sal group. # p < 0.05 and ## p < 0.01 compared with Control + LPS group using the one-way ANOVA with Bonferroni’s post hoc test or or Kruskal-Wallis test followed by Dunn’s post hoc test. Control + Sal (*n* = 8-12), Control + LPS (*n* = 9-10), SAD + Sal (*n* = 5), and SAD + LPS (*n* = 7-13).

#### Spleen

Spleen is considered the efferent component of the “splenic anti-inflammatory reflex” (20). In the spleen, we observed a significant increase in TNF-α (P = 0.0242), IL-6 (P = 0.0026), IL-10 (P < 0.0001), and IFN-γ (P = 0.05) in Control + LPS in comparison with Control + Sal rats. In addition, we observed reduced surges of TNF-α (P = 0.7554) or IFN-γ levels (P > 0.9999) in SAD + LPS group in relation to Control + LPS animals (Fig. 8 E, F, G and H). These findings indicate that SAD reduces inflammatory signaling in spleen during SI.

### Corticosterone, NOx, and norepinephrine during LPS-induced SI

Plasma corticosterone levels were increased in Control + LPS (P < 0.0001) and in SAD + LPS (P < 0.0001) in comparison with Control + Sal group. There was no significant difference between Control + LPS and SAD + LPS groups (P > 0.9999, Fig. 9 A). Interestingly, the observed LPS-induced drop in MAP was accompanied by increased plasma nitrate concentration, and these changes were independent of baroreceptor afferents integrity (P < 0.0001, Fig. 9 B). Furthermore, plasma norepinephrine levels were similar in all the evaluated groups (Fig. 9 C, P = 0.7053).

**Fig. 9:**
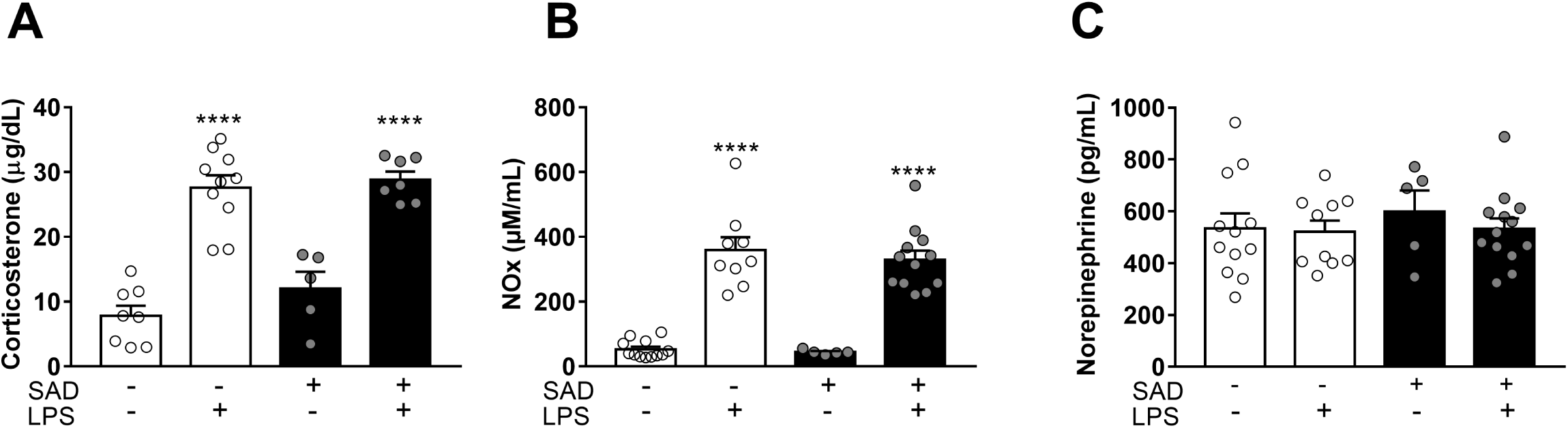
Plasma levels of corticosterone (panel A), NOx (nitrate, panel B), and norepinephrine (panel C) from Control + Sal, Control + LPS, SAD + Sal and SAD + LPS groups. Results are presented as individual values and mean ± SEM. **** p < 0.0001 compared with Control + Sal group using the one-way ANOVA with Bonferroni’s post hoc test. Control + Sal (*n* = 9), Control + LPS (*n* = 8-9), SAD + Sal (*n* = 5-6), and SAD + LPS (*n* = 5-9).

## Discussion

The present study is the first to report that baroreceptors afferents are key to modulate not only LPS-induced SI but also its consequent cardiovascular changes and to provide evidence that these phenomena are not dependent of each other. Supporting this notion, we observed that the LPS-induced hypotension was higher and earlier in rats previously submitted to SAD than in control rats. We also show that HF power of PI was altered in rats that received LPS, while LPS-induced increase in LF component of SAP was dependent on sino-aortic afferents integrity, even in the presence of vascular dysfunction. Of particular importance, we observed reduced surges of plasma interleukin (IL)-6 and IL-10 and splenic TNF-α and interferon-γ in SAD rats, indicating that SAD not only modulated LPS-induced cardiovascular collapse but also reduced peripheral cytokines surges. Moreover, this reduced LPS-induced plasma cytokines surges seems to be at least in part mediated by the splenic anti-inflammatory reflex (20), since reduced levels of pro-inflammatory cytokines in the spleen were observed after SAD (Fig. 8).

### Cardiovascular control during SI is dependent on baroreceptor afferents integrity

Considering that baroreceptor afferents integrity is important in the moment-to-moment control of cardiovascular function (21), we recorded blood pressure and HR in conscious rats through 3 hours after LPS administration. LPS-induced hypotension was significantly enhanced after SAD (Fig. 1). In contrast, LPS-induced tachycardia was blunted in SAD rats. Tachycardia during SI is a compensatory mechanism regulating hypotension (17) which is associated with vascular dysfunction (22). Blunted tachycardia with greater hypotension in SAD rats may be related, at least in part, to a baroreflex-dependent cardiovascular regulation during SI. Several studies document that baroreflex modulates sympathetic and parasympathetic activity to the heart and resistance blood vessels in health and disease, including SI (11,23,24). Our results are consistent with the notion that the LPS-induced hypotension (Fig. 1), caused by a reduction in vascular resistance (Fig. 6 and 7) even in the presence of increase of sympathetic vasomotor tone (Fig. 3), is critically dependent upon the baroreceptors afferents integrity in conscious rats. These data are in agreement with prior studies showing a significant fall in blood pressure of anesthetized rats that received a lethal dose of LPS (15 mg.kg^−1^, given IP) and in which carotid chemo and baroreceptors were previously denervated (25). Moreover, it has been reported that SAD rats show a significant reduction in the survival time during polymicrobial sepsis, indicating that baroreflex dysfunction is associated with a poor sepsis prognosis (26). Of special significance, LPS-induced tachycardia was significantly blunted in SAD + LPS rats, indicating that in our experimental condition, baroreflex integrity is involved, at least in part, with increased HR during SI.

Regarding HR variability in the time-domain, LPS-induced SI acutely reduced SDNN and in RMSSD, which were not affected by SAD (Fig. 2). These findings are consistent with the notion that during SI, an imbalance in the autonomic cardiac control takes place. Previous studies indicate that both LPS or TNF-α administration significantly reduces HR variability in mice indicating a causal link between cytokines and abnormal SDNN (27). More recently, clinical studies point out that HR variability is significantly increased in athletes and reduced in patients (28). In respect of SAP variability, the SD of MAP was only sustained in SAD + Sal group, suggesting that the hallmark of SAD experimental model, [higher variability of the MAP (13)] is critically affected during SI.

In the search for mechanisms underlying hemodynamic control during SI, we used spectral analysis of PI in the frequency domain, which showed that HF component of PI was significantly reduced at 3 hours after endotoxin in Control + LPS and in SAD + LPS in relation to Control + Sal (Fig 3 A and D). These observations are in line with the concept that LPS-induced tachycardia is triggered by mechanisms resulting in a decrease of vagal modulation to the heart (18). Significant changes in the autonomic control to the heart have been observed during the initial phase of sepsis-associated SI as a compensatory adjustment to avoid circulatory shock (29). Regarding the variance of SAP in the frequency domain, the LF component was significantly increased in Control + LPS rats than in Control + Sal and SAD + LPS (Fig. 3), indicating that LPS-induced increases in sympathetic vasomotor tone depends on sino-aortic afferents integrity even in the presence of vascular dysfunction. This eventual sustained sympatho-excitation during SI is in agreement with a previous study (17) and suggest that sympathetic drive to resistance vessels does not revert hypotension in this experimental model.

In addition to linear methods (time and frequency-domain analyses), non-linear approaches were also used in the present study. Methods for analysis of nonlinear dynamics has been utilized to increase the interpretation of the complexity cardiovascular function (30, 31). We found that DFA α_2 scaling exponent of SAP was reduced in Control + LPS rats and was greater in SAD + Sal and in SAD + LPS than in Control + Sal rats (Fig. 4). These findings are consistent with the notion that the control of blood pressure is highly complex and that during LPS-induced SI the oscillations in SAP tend to erratic or random patterns (α_2 < 1). In contrast, when LPS is administrated to SAD animals, SAP oscillations tend to be smoother (α_2 > 1). Reconciling the data obtained from MSE curves, we observed that SAP entropy of the SAD + Sal and Control + LPS was significantly different from those for Control + Sal group (Fig. 5), suggesting the that the baroreflex plays an important role in the complex response to challenges imposed to the cardiovascular system (30). Although a direct interpretation of the functional meaning of these results is not an easy task, the analyses of SAP complexity has been used to predict cardiovascular outcomes and has been associated with high mortality risks (32). As far as we know, this is the first study that analyzed these cardiovascular complexity patterns by nonlinear approaches in SAD rats during SI. Whether or not SAD-induced changes in cardiovascular complexity may worse hemodynamic collapse during SI is an interesting matter deserving further investigation.

### Vascular dysfunction during SI or SAD are not cumulative

Considering previous studies (17,18,33) and our own data (Fig. 1) documenting cardiovascular collapse during SI, we further evaluated the vascular function after LPS administration in Control and SAD animals in mesenteric resistance arteries. After LPS administration, a significant reduction in contraction induced by Phe and by EFS in mesenteric arteries from LPS-treated Control and SAD groups occurs (Fig. 6 and 7). Vascular hyporesponsiveness was reverted by the nitric oxide synthase inhibitor L-NAME, indicating that LPS-induced SI reduces contraction of resistance arteries by mechanisms that involve nitric oxide synthase activation whereas may be independent of baroreceptors afferents integrity. It is important to point out that LF component of SAP (an index of sympathetic vasomotor tone) was observed to be increased in Control + LPS conscious rats and that vascular responsiveness to Phe was observed to be decreased in mesenteric resistance arteries of these animals in comparison with Control + Sal suggesting that in SI changes in the sympathetic nerves modulation to alpha1-adrenergic receptors in mesenteric resistance arteries take place.

Additionally, the endothelium-dependent relaxation response induced by cumulative concentrations of ACh was similar between the all Control and SAD groups groups (Fig. 6), indicating that neither SAD itself nor LPS administration cause endothelium dysfunction in our experimental condition. Conversely, endothelium-independent relaxation response was reduced in mesenteric arteries from all the groups that received LPS, probably due to changes in the sensitivity to guanylate cyclase/ cGMP pathway (34). The hypothesis that the guanylate cyclase/cGMP pathway is affected during SI is possible given that NO-induced activation of guanylate cyclase enzyme in vascular smooth muscle cells leads to recruitment of intracellular signaling cascades that reduce intracellular Ca^2+^ levels, open K^+^ channels and cause relaxation (35).

### Changes in Tb during SI are affected by SAD

Interestingly, rats submitted to SAD presented no fever, but a significant fall in Tb (hypothermia) after LPS administration (Fig. 1). The mechanisms involved in LPS-induced hypothermia are not completely known. Nevertheless, SI-associated hypothermia has considerable clinical implications (36). One may consider LPS-induced hypothermia as a failure in neural control of Tb. Alternatively, recent studies have suggested that hypothermia is precisely controlled by specific mechanisms mediated by the central nervous system (37). Based on these data, we suggest that baroreceptor afferents integrity affect thermoregulatory control during SI by impairing a key part of the afferent signals to the brain. We further speculate that among the important multimodal functions of arterial baroreceptors (21) febrigenic signaling in the periphery affects brain circuitry at least in part by interacting with peripheral baroreceptors afferents. The reduced LPS-induced plasma surges of IL-6 in SAD rats (Fig. 8) may be at least one of the contributing factors in thermogenesis during SI, since this cytokine is known to induces increases in Tb acting as an important endogenous pyrogen (38). These baroreceptor afferents effects on thermoregulation must take place through thermoeffectors modulation, but it remains unknown if this modulation is via sympathetic innervation of the brown adipose tissue (affecting non-shivering thermogenesis) or sympathetic innervation of the tail artery (affecting heat loss index).

### Plasma and spleen surges of pro-inflammatory cytokines during SI are affected by SAD

The LPS-induced plasma surges of IL-6 (a pro-inflammatory cytokine) and IL-10 (an anti-inflammatory cytokine) in SAD rats were significantly reduced after SAD (Fig. 6). These findings indicate that baroreflex does modulate the LPS-induced peripheral cytokine surges and adds new information to a previous study that documented that electrical baroreflex stimulation inhibits LPS-induced pro-inflammatory cytokines surges not in the periphery but in the brain (19). A putative mechanism by which sino-aortic afferents may play a modulatory effect in the LPS-induced IL-6 and IL-10 plasma surges may be related to macrophages polarization, a highly heterogeneous cell population, during SI. Activated macrophages M1 exhibit high levels of pro-inflammatory cytokines, while activated M2 macrophages exhibit high levels of anti-inflammatory cytokines (39).

In addition to circulating macrophages aforementioned, the main efferent target organ for the splenic anti-inflammatory reflex is the splenic macrophages located in the white pulp (2). We showed here that spleen tissue homogenates collected from rats that received LPS exhibit a significant increase of pro-inflammatory cytokines. Surprisingly, there were no significant increases in TNF-α [an essential early mediator of inflammation (2, 40)] or IFN-γ [a later inflammatory marker (41)] levels in rats submitted to SAD during LPS-induced SI (Fig. 8 B). Combining a previous study showing that stimulation of the efferent fibers impinges upon the spleen leads to a significant anti-inflammatory effects in this organ during LPS-induced SI (42) and our own data in which SAD rats showed decreased LPS-induced surges of pro-inflammatory cytokines in the spleen, we suggest that the efferent arm of the splenic anti-inflammatory reflex is modulated by baroreceptors afferents.

Considering that baroreflex stimulation downregulates pro-inflammatory cytokines in hypothalamus, but not in plasma, heart and spleen (19) we hypothesized that SAD worsens cytokines surges in plasma. Contrary to our expectations, after SAD, LPS-induced surges of cytokines were blunted in plasma and spleen (Fig 8.) suggesting that the baroreceptor afferents integrity/stimulation may differentially affect peripheral and central pro-inflammatory cytokines surges in this critical condition that resembles some features of sepsis, a considerable healthcare burden.

A plethora of studies have provided strong evidence demonstrating autonomic regulation of immune function (2, 43). For instance an inhibitory action of the sympathetic nervous system and its main neurotransmitter, norepinephrine on SI has been documented (2, 43). In the present study, we show that the known LPS-induced enhancement of sympathetic vasomotor tone (23, 43) may be attributable, at least in part, to sino-aortic afferents integrity (Fig. 3). Whether the LPS-induced sustained increase in the activity of the splenic nerves (2) is affected by sino-aortic afferents is a possibility that requires additional investigation. In addition, if pro-inflammatory cytokines surges in SI (2) are regulated by baroreceptor afferents or if this exacerbate release of immune mediators represents a failure of the adaptive mechanisms are interesting matters to be explored.

Even though we observed that SAD affects LPS-induced cardiovascular collapse, thermoregulation and inflammatory signaling, we can make no conclusions about the causal link between these important regulatory functions, reflecting the complexity of this experimental model in which a myriad of events lead to multiple organ failure and eventually death depending on the doses of LPS. We suggest that the baroreflex-dependent mechanisms mediating inflammatory status are not associated with cardiovascular collapse, given that SAD reduced cytokines surges (both in plasma and spleen) and exacerbate hypotension.

### Corticosterone, NOx and norepinephrine during SI are not affected by SAD

In SI, a significant increase in plasma corticosterone (a hormone with anti-inflammatory action) levels occurs. This LPS-induced increased corticosterone levels were not affected in SAD rats (Fig. 9 A). These data support the notion that during LPS-induced SI an increase in the hypothalamic-pituitary-adrenal axis activity occurs (44) and that this activation is independent of the baroreceptor afferents integrity.

To provide insights into the mechanisms involved in hypotension during SI, we also assessed plasma NO (a potent vasodilator) and norepinephrine (a vasopressor neurotransmitter) levels (Figs 9). Taking into consideration that during sepsis, NO pathway system is markedly stimulated leading to decreased vascular responsiveness to constrictor stimuli (22), our findings further support the notion that indeed LPS-induced SI is accompanied by a significant increase in plasma NO production, that does not depend on baroreceptors integrity. Moreover, these findings indicate that greater hypotension in SAD rats is not associated with higher NO production, but rather to the autonomic imbalance *per se* (Fig. 9 B). Siminarly, the observed effects of SAD on the LPS-induced hemodynamic dysfunction seems to be independent of systemic noradrenaline levels, since this catecholamine levels were similar among groups (Fig. 9 C). However, these data do not rule out that local noradrenaline release from sympathetic nerve terminals during SI may be different depending on the vascular bed.

In conclusion, the present data are consistent with the notion that the role of baroreflex afferents on LPS-induced SI goes beyond the lessening hypotension and tachycardia despite severe vascular dysfunction and affecting inflammatory status. The present findings shed light on the mechanisms underlying the contribution of cardiovascular afferents in the regulation of the inflammatory surges in plasma and spleen during SI.

## Materials and Methods

All animal experimentation was executed according to directions for animal study from the National Council for Animal Experimentation Control in Brazil (CONCEA). The experimental procedures were also reviewed and approved by The Ethics Committee on Animal Research of the Dental School of Ribeirão Preto - University of São Paulo, Ribeirão Preto, Brazil (#2017.1.585.58.9).

### Animals

Male Wistar rats (300–350 g) were acquired from the Animal Care Facility of the University of São Paulo at Ribeirão Preto. During experiments they were kept in plastic cages in the animal facility of the Dental School of Ribeirão Preto, University of São Paulo under a 12-h light/dark cycle (lights on at 6 am) at 23-24 °C. Rats had unrestricted access to standard chow and tap water.

### Surgical procedures

#### Sino-aortic denervation

The most well accepted model of baroreceptor afferents removal, called sino-aortic denervation (SAD) (45,23,46), was performed aseptically using a standard technique (47). Briefly, rats were anesthetized with a cocktail of ketamine (100 mg.kg^−1^) and xylazine (10 mg.kg^−1^) and fixed in the supine position after the absence of the withdrawal reflex to tail and paw pinch. Additional doses of anesthetic were administrated if necessary. A ventral midcervical incision was performed and fibers from the aortic depressor nerve traveling with the superior laryngeal nerve and superior cervical ganglion were transected. The carotid baroreceptors were denervated by removal of surrounding tissues from the carotid sinus.

#### Arterial and venous catheterization

On the fourth day after SAD rats were anaesthetized with ketamine and xylazine and a polyethylene catheter (PE-10 connected to PE-50 tubing; Clay Adams, Parsippany, NJ, USA, Intramedic, Becton Dickinson, Sparks, MD, EUA), was placed into the abdominal aorta by means of femoral artery. Femoral vein was also catheterized. Artery catheterization was used to direct hemodynamic recordings while vein catheter was utilized for drug administration. Both catheters were tunneled subcutaneously and exteriorized through the skin in the nape of the neck, and the surgical wounds were sutured aseptically. Rats recovered individually in the recording room. On the following day the arterial catheter was connected to a pressure transducer (MLT0380; ADInstruments), and in turn, to an amplifier (Bridge Amp, ML221; ADInstruments). Pulsatile arterial pressure (PAP) and heart rate (HR) were recorded using the Chart Pro software (ADInstruments) were recorded simultaneously placed in side-by-side cages. Rats from different groups were recorded simultaneously placed in side-by-side cages. Beat-by-beat series of systolic arterial pressure (SAP) and PI were obtained from the raw PAP recordings and SAP or PI variability was evaluated using the software CardioSeries (48) and JBioS (49).

In the time domain, standard deviation (SDNN) and root mean square of the successive differences (RMSSD) were calculated from PI series. Standard deviation (SD) was also obtained from SAP series. In the frequency domain, the power spectra of PI and SAP were estimated by the modified periodogram and Welch protocol (50). Briefly, all series were interpolated at 10 Hz (cubic spline) and divided into segments of 512 points (51.2 seconds). Segments containing artifacts or transients were excluded. Next, each selected segment was multiplied by a Hanning window and the periodogram was estimated. The PI spectra were integrated into low- (LF, 0.2–0.75 Hz) and high-frequency (HF, 0.75–3 Hz) bands, while the SAP spectra were integrated at LF band only. The power at LF band was assessed in normalized units (nu), represented by LF/(LF+HF), whereas the power at HF band was evaluated in absolute units. According to previous studies (31, 51), this representation provides the best correlation of spectral indices to the sympathetic and parasympathetic modulation of the heart rate, respectively.

Nonlinear properties of PI and SAP series were assessed by multiscale entropy (MSE) and detrended fluctuation analysis (DFA) (52, 53). MSE quantifies the degree of irregularity (unpredictability) of time series over increasing time scales and can be considered a measure of physiological complexity. Healthy systems represent the most complex physiological status, whereas aging and diseases denote some disruption in the integrative regulatory mechanisms, decreasing the capability of the organism to adapt to changing demands (54). MSE parameters were set to m=2 (embedding dimension), r=15% of time series SD (tolerance factor) and τ=1…20 (time scales). On the other hand, DFA quantifies the power law scaling of time series, which is related to its fractal temporal structure (55). In the present study, α_1 comprises windows from 5 to 15 points and α_2 comprises windows from 30 to 10 and α_3 comprises windows from 100 to N⁄10 points, where N is the time series length.

#### Temperature datalogger implantation

In the same surgical procedure for arterial and venous catheterization, a median laparotomy was done and an intraperitoneal temperature data-logger capsule (SubCue, Calgary, AB, Canada) was inserted to deep body (Tb) temperature recordings in rats. Afterward, surgical wounds were sutured aseptically.

### Vascular reactivity studies

The method described by Mulvany and Halpern (1977) (56) was used. Animals were euthanized and segments of third-branch mesenteric arteries, measuring about 2 mm in length, were mounted in a small vessel myograph (Danish Myo Tech, Model 620M, A/S, Århus, Denmark). Arteries were maintained in a Krebs Henseleit solution [(in mM) NaCl 130, KCl 4.7, KH2PO4 1.18, MgSO4 1.17, NaHCO3 14.9, Glucose 5.5, EDTA 0.03, CaCl2 1.6], at a constant temperature of 37 °C, pH 7.4, and gassed with a mixture of 95% O2 and 5% CO2.

Mesenteric resistance arteries were set to reach a tension of 13.3 kPa (kilopascal) and remained at rest for 30 min for stabilization. The arteries were stimulated with Krebs solution containing a high concentration of potassium [K^+^, (120 mM)] to evaluate the contractile capacity of the segments. After washing and return to the basal tension, arteries were contracted with Phe (10^−6^ M) and then stimulated with ACh (10^−5^ M) to determine the presence of a functional endothelium. Arteries exhibiting a vasodilator response to ACh greater than 80% were considered endothelium-intact vessels. After washing and another period of stabilization, concentration-response curves to Phe (10^−10^ to 3×10^−5^) and EFS were performed in mesenteric resistance arteries to produce contractions, measured as increases in baseline tension. EFS was applied to arteries placed between platinum pin electrodes and conducted at 20 V, 1-ms pulse width, and trains of stimuli lasting 10 s at varying frequencies (1 to 32 Hz).

Vasodilation responses were determined in mesenteric resistance arteries contracted with Phe (10^−6^ to 3×10^−6^ M). After 15 min, concentration-response curves to ACh (10^−10^ to 3×10^−5^ M) and SNP (10^−10^ to 10^−5^ M) were carried out. Concentration-response curves to Phe, EFS and ACh were also performed in the presence of L-NAME (10^−4^ M).

### Plasma measurements

At the end of the cardiovascular and Tb recordings, arterial blood was withdrawn in EDTA-coated tubes and centrifuged (20 min at 3.500 rpm, 4 °C), for plasma extraction, on the third hour after saline or LPS administration. All plasma samples were kept at −80 °C until assays.

#### Cytokines and noradrenaline

Plasma samples were assayed for measurement of tumor necrosis factor (TNF)-α, interleukin (IL)-6, IL-10 and interferon (IFN)-γ using multiplex assay kits according to standard instructions (LXSARM - 05, R&D System, Minnesota, USA) with Luminex® Magpix™ technology (Austin, TX, USA). Plasma samples were assayed for measurement of noradrenaline (Cloud-Clone, Texas – USA) levels, using enzyme-linked immunosorbent assay (ELISA) kits according to standard instructions.

Spleens were homogenized in 0.5 mL of PBS, protease inhibitor cocktail (Cell Signaling, Massachusetts, USA) and then centrifuged at 13,000 rpm for 20 min at 4 °C. Tissue supernatant samples were used to measure TNF-α, IL-6, IL-10, and IFN-γ levels by a multiplex assay as in plasma samples. Data from splenic cytokines were normalized by protein concentrations by means of Bradford assay (#5000205, Bio-Rad Laboratories, USA).

#### Radioimmunoassay for corticosterone

Plasmatic corticosterone extractions and radioimmunoassay were performed from 25 μL of plasma by adding 1 mL of ethanol according to Haack et al., (1979) (57).

#### Nitric oxide (nitrate, NOx)

Plasma NOx levels were assessed by using the chemiluminescence NO-Ozone technique. Nitrate concentrations were measured using 40 μL aliquots of the plasma samples inserted into a NO analyzer (model 280, Sievers Instruments, Boulder, CO, USA).

### Experimental Design

Rats were assigned into 4 experimental groups:

Control + Sal: Naïve rats that received saline administration.

Control + LPS: Naïve rats that received LPS administration.

SAD + Sal: SAD rats that received saline administration.

SAD + LPS: SAD rats that received LPS administration.

#### 1) Protocol #1

To study the role of sino-aortic afferents integrity in LPS-induced changes in MAP, HR and Tb, rats were catheterized and had a datalogger implant and on the day after they received an iv injection of saline or LPS and were recorded up to 180 min after iv administration. To avoid the influence of variability of MAP of SAD rats in the results, the reported values of MAP and HR were obtained as the delta of beseline values (using a mean of 10 min of recording before the intravenous injection) and the minimum value for MAP and the maximum value for HR obtained from the last minute of every 10 min period throughout 180 min. This analysis was done in all the experimental groups.

#### 2) Protocol #2

To further characterize the role of sino-aortic afferents integrity in LPS-induced cardiovascular collapse, spectral analysis of PI and SAP in the time and in the frequency domain, and analyze of the complexity of cardiovascular function were evaluated offline.

#### 3) Protocol #3

180 min after LPS or saline administration, arterial plasma was withdrawn to assess corticosterone, NOx, and norepinephrine levels. Cytokines levels were also assessed in plasma and spleen.

#### 4) Protocol #4

180 min after LPS or saline administration, rats were euthanized and vascular reactivity was evaluated in mesenteric resistance arteries.

### Statistical analysis

Data are expressed as mean ± S.E.M. (standard error of the mean) and significant differences were considered at P ≤ 0.05, but exact P values are described. Unpaired t test, one-, two-way ANOVA followed by the Bonferroni multiple comparisons test or Kruskal-Wallis test followed by Dunn’s multiple comparisons test were performed when necessary.

## Conflict of interest

The authors declare no conflict of interest.

## Acknowledgments

We are grateful to Maria Valci dos Santos and Mauro F. Silva for their excellent technical assistance.

## Funding

This work was supported by Grant L.G.S.B [#2016/17681-9, São Paulo Research Foundation (FAPESP)], fellowship to M.R.A. [#2017/09878-0 São Paulo Research Foundation (FAPESP)] and National Council for Scientific and Technological Development (CNPq), Brazil.

## Author Contributions

M.R.A., J.L.D., C.A.P., G.S.B. and N.S.F. performed experiments. M.R.A., L.E.V.S. and C.A.P., analyzed the data. M.R.A., J.L.D. and L.G.S.B conceived and designed the study. M.R.A., and L.G.S.B. planned the experiments, wrote, reviewed and contributed to the final manuscript. L.G.S.B., J.A.R., E.C.C., and R.C.T. supervised the project and provided funding. All authors reviewed the manuscript

